# Updated benchmarking of variant effect predictors using deep mutational scanning

**DOI:** 10.1101/2022.11.19.517196

**Authors:** Benjamin J. Livesey, Joseph A. Marsh

**Affiliations:** MRC Human Genetics Unit, Institute of Genetics and Cancer, University of Edinburgh, Edinburgh EH4 2XU, UK

## Abstract

Variant effect predictors (VEPs) provide a potential solution to the influx of variants of uncertain clinical significance produced by genome sequencing studies. However, the assessment of VEP performance is fraught with biases introduced by benchmarking against clinical observations. In this study, building on our previous work, we use independently generated measurements of protein function from deep mutational scanning (DMS) experiments for 26 human proteins to benchmark 55 different VEPs, while introducing minimum data circularity. The top VEPs are dominated by unsupervised methods including EVE, DeepSequence and ESM-1v, a new protein language model that ranked first overall. However, the strong performance of recent supervised VEPs, in particular VARITY, shows that developers are taking data circularity and bias issues seriously. We also assess the performance of DMS and unsupervised VEPs for discriminating between known pathogenic and putatively benign missense variants. Our findings are mixed, demonstrating that some DMS datasets perform exceptionally at variant classification, while others are poor. Notably, we observe a striking correlation between VEP agreement with DMS data and performance in identifying clinically relevant variants, with EVE, DeepSequence and ESM-1v performing best, further supporting the utility of DMS as an independent benchmark.

## Introduction

Accurately classifying variants of uncertain clinical significance remains an ongoing challenge for variant interpretation. The most common type of genetic variant in humans are single nucleotide variants, most of which have no role in disease (Auton *et al*, 2015). Pathogenic variants are enriched among the rarest occurring variants in the human population (Wang *et al*, 2021), which makes gathering sufficient evidence to classify them challenging. Over the past two decades the field of computational variant effect prediction has striven to provide additional evidence for the classification of variants of uncertain significance frequently identified in genetic sequencing data (Livesey & Marsh, 2022). Variant effect predictors (VEPs) are algorithms that use evidence from various sources, including evolutionary conservation, functional annotations, and physicochemical differences, to predict the likely phenotypic outcome of a genetic variant. The output of VEPs must be benchmarked against a “gold standard” to ensure that the predictor is generating accurate results (Sarkar *et al*, 2020). Such benchmarking studies are frequently conducted both by VEP authors and independent groups, traditionally by comparing VEP classifications to sets of known pathogenic and benign variants (Gunning *et al*, 2021; Niroula & Vihinen, 2019). This approach raised some concern over the potential for data circularity (the reuse of data) to inflate VEP performance estimates (Grimm *et al*, 2015). Type 1 circularity involves re-using data originally used to train a predictor while assessing its performance, leading to improved performance compared to more appropriate benchmarking data. Type 2 circularity occurs when a VEP identifies a gene where mutations are highly skewed towards either a pathogenic or a benign outcome. In these cases, future predictions on mutations in this gene may be influenced by a VEPs previous experience, often resulting in good performance in other mutations in these proteins, but much poorer performance on novel proteins or genes with mixed clinical outcomes associated with mutations.

We previously attempted to address the issue of data circularity by using data from deep mutational scanning (DMS) studies as the “gold standard” to perform a benchmark of VEP performance against single amino acid variants (Livesey & Marsh, 2020). DMS encompasses a wide variety of high-throughput experimental techniques, whereby functional scores for large numbers of amino acid variants are measured (Fowler & Fields, 2014). Because most DMS-derived functional scores are for variants never observed in the human population, using them to assess VEP performance can address the issues of limited benchmarking data availability that sometimes lead to type 1 circularity. Even DMS-derived variants that exist in VEP training data have functional scores fully independent from previous clinical labels. Our study also used the correlation between the continuous outcome of each VEP and the DMS functional scores as the basis for our benchmark. This approach helps to address type 2 circularity as a VEP cannot score highly by assigning all variants in a protein as a single class, but must determine the relative functional impact of each variant. Previously, we identified a method based on unsupervised machine learning, DeepSequence (Riesselman *et al*, 2018), to be the top-performing VEP for human proteins. We also demonstrated the ability of DMS to outperform VEPs at direct classification of clinically relevant variants.

Significant progress has been made in both VEP development and DMS methodologies since our previous study with multiple predictors based on cutting edge machine learning methodology and many new DMS studies being published. In this paper, we have updated our previous benchmarking strategy with the addition of more recently published VEPs, and many additional human DMS datasets. We also use known pathogenic missense variants to assess the performance of DMS and unsupervised VEPs for predicting the outcome of known missense variants. While DMS has huge potential as a benchmarking tool and in direct variant classification, the results of DMS assays do not necessarily correlate with human disease outcomes. One must ensure that the fitness metric being measured is relevant to human pathogenic conditions. It is also not currently feasible to perform DMS assays for the entire human proteome, so VEPs will likely play a significant role in variant interpretation for some time to come.

## Results

### Overview of VEPs and DMS datasets used in this study

Compared to our previous benchmark, we increased the number of DMS datasets of human single amino acid variants to 26 and the total number of VEPs, conservation scores and substitution matrices benchmarked to 55. We considered exclusively human proteins, as only a subset of the VEPs we include in this analysis can generate predictions for non-human proteins.

We identified new and previously unused DMS datasets through searching MaveDB (Esposito *et al*, 2019) and identifying recently published works in the literature. Table 1 summarises each of the new DMS studies that were added to the analysis, with the full set of DMS experiments given in Table S1.

**Table 1.** DMS assays included in this study that were not present in our previous benchmark.

| **DMS target**<br>**(Uniprot ID)** | **Functional assay** | **Coverage of all**<br>**amino acid**<br>**substitutions (%)** | **Reference or**<br>**MAVEDB**<br>**identifier** |
| --- | --- | --- | --- |
| **SNCA** (P37840) | Yeast growth rate hindered by aggregate toxicity (reverse-survival) | 97.26 | (Newberry *et al*, 2020) |
| **CASP3**<br>(P42574)<br>**CASP7**<br>(P55310) | Apoptotic activity assessed by fluorescence in a microfluidic system. | 28.63<br>29.17 | (Roychowdhury & Romero, 2022) |
| **CBS** (P35520) | Yeast growth rate complementation | 64.41 | (Sun *et al*, 2020) |
| **CCR5** (P51681)<br>**CXCR4**<br>(P61073) | Antibody binding activity and surface expression levels in human cells | 99.97<br>99.36 | (Heredia *et al*, 2018) |
| **CYP2C9**<br>(P11717) | Activity profiling (Click-seq). | 65.97 | (Amorosi *et al*, 2021) |
| **GDI1** (P31150) | Yeast growth rate complementation | 51.40 | (Silverstein *et al*, 2021) |
| **HMGCR**<br>(P04035) | Yeast growth rate complementation | 99.89 | (Jiang, 2019) |
| **LDLRAP1**<br>(Q5SW96) | Yeast two hybrid binding assay | 99.03 | (Jiang, 2019) |
| **MSH2** (P43246) | Rescue of MMR-deficient HAP1 cells | 94.38 | (Jia *et al*, 2021) |
| **MTHFR**<br>(P42898) | Yeast growth rate complementation | 99.85 | (Weile *et al*, 2021) |
| **NUDT15**<br>(Q9NV35) | Drug resistance assay (growth rate). | 94.16 | (Suiter *et al*, 2020) |
| **TP53** (P04637)<sup>1</sup> | reverse growth rate assay in human cells | 39.37 | (Kotler *et al*, 2018) |
| **PDE3A**<br>(Q14432) | DNMDP sensitivity in a glioblastoma cell line. | 36.41 | (Garvie *et al*, 2021) |
| **VKORC1**<br>(Q9BQB6) | Protein stability assessed by FACS (VAMP-seq). | 87.02 | (Chiasson *et al*, 2020) |

<sup>1</sup> - Our previous benchmark already included TP53, but we identified a further dataset published by another group.

Many DMS datasets provide multiple scores covering different experimental conditions and sometimes entirely different fitness assays of the same protein; mappings between the names of these assays and identifiers used in this study are provided in Table S2. We calculated the absolute Spearman’s correlations between these score sets in the same protein to gauge the reproducibility of DMS results under different conditions. The strongest correlations (>0.9) were between biological replicates, while assays investigating fitness under different experimental conditions or alternate fitness metrics often resulted in much lower correlations (<0.3). Most correlations observed between alternative assays were in a range between 0.4 and 0.6 (Table S3), which is similar to the level of correlation between DMS and the top VEPs in our previous study. To represent each protein in our analysis, we selected a single assay from each DMS study. For proteins with multiple DMS datasets available, the assay that gave the highest median absolute Spearman’s correlation against all VEPs was selected to be representative of fitness effects in each DMS target protein (Table S1). The use of the median ensures that our assay selection is not skewed by a few particularly high or low-correlating VEPs.

Several of the new VEPs included in this analysis were added to the dbNSFP database in the 4.2 update (Liu *et al*, 2020), while others were identified by literature search. We also removed VEPs that were no longer accessible (NetDiseaseSNP) or unsuitable for a benchmark against continuous data (MutationTaster). A summary of the new VEPs assessed in our benchmark along with their sources is provided in Table 2, while the full list of all VEPs is in Table S4.

**Table 2.**
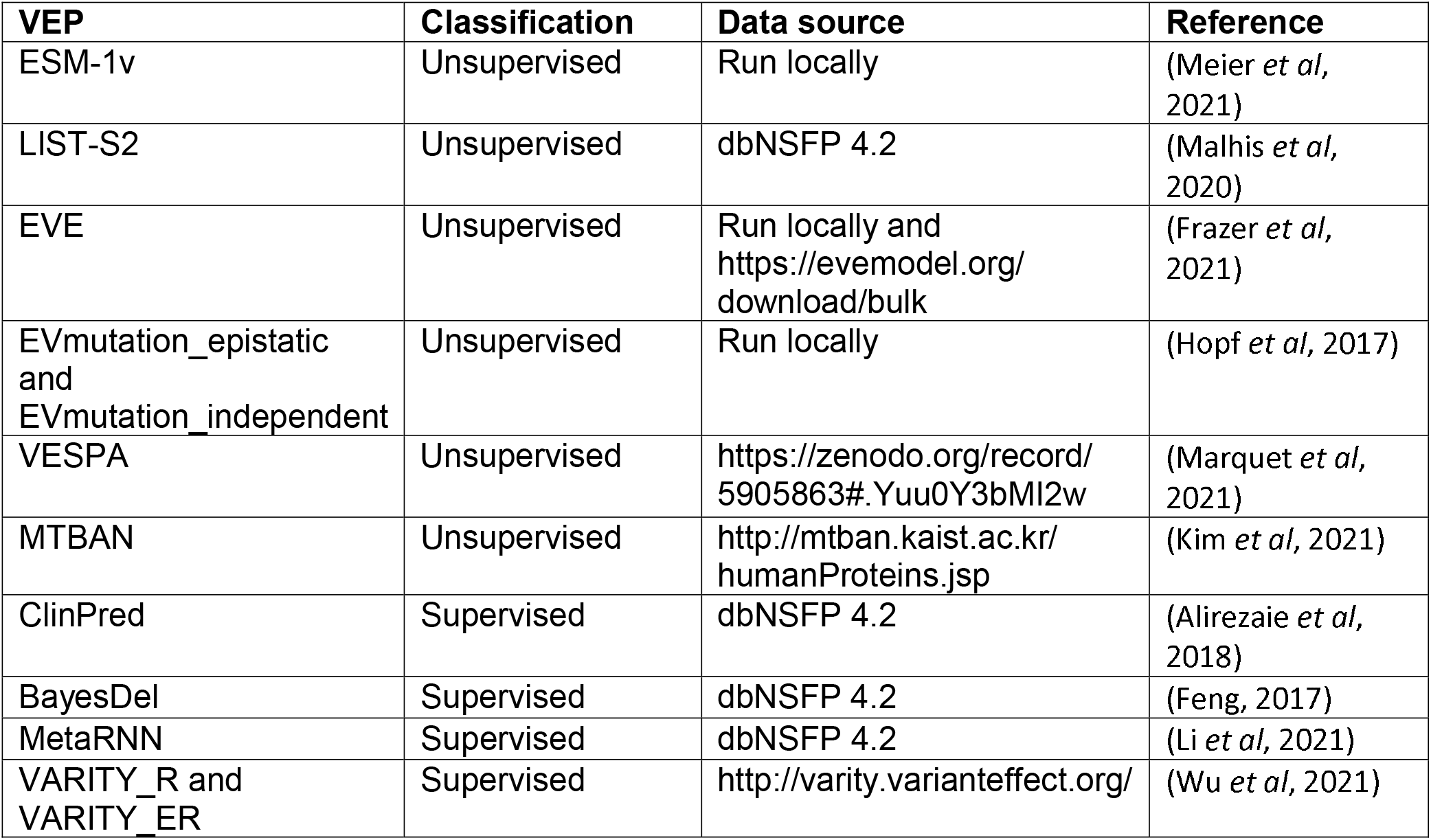
VEPs benchmarked in this study that were not present in our previous results.

Previously, we defined four different categories to classify VEPs based on their architecture and training: supervised, unsupervised, empirical and metapredictors. These categories overlapped with each other to some extent as several VEPs could fall into multiple categories. To better reflect which predictors are related by methodology we have now given all VEPs a label that is either “supervised” or “unsupervised” (Table 2, Table S4), which reflects whether labelled examples were used to train the predictor and thus whether data circularity is a concern for its assessment. Despite this simplification of VEP classification, Eigen could still qualify for both categories. Eigen uses an unsupervised spectral method to combine multiple other VEP scores and deleteriousness metrics. However, one of the VEPs it includes as a feature is PolyPhen-2, a supervised VEP that may act as an indirect source of data circularity. Thus, Eigen has the potential for data circularity, and we have therefore labelled it as supervised in this analysis.

### Benchmarking of VEPs using DMS data

We calculated the Spearman’s correlation between each of the selected representative DMS datasets for every protein, and all available variant effect predictions using the continuous outcome scores produced by each VEP. Our results show that many of the recently developed VEPs produce superior performance to those already present in our previous analysis (Fig 1). Of particular note are the unsupervised methods EVE and ESM-1v as well as the supervised predictor VARITY. EVmutation is a slightly older unsupervised VEP that was not included in our previous study, but also produced high correlations with DMS data.

**Figure 1.**
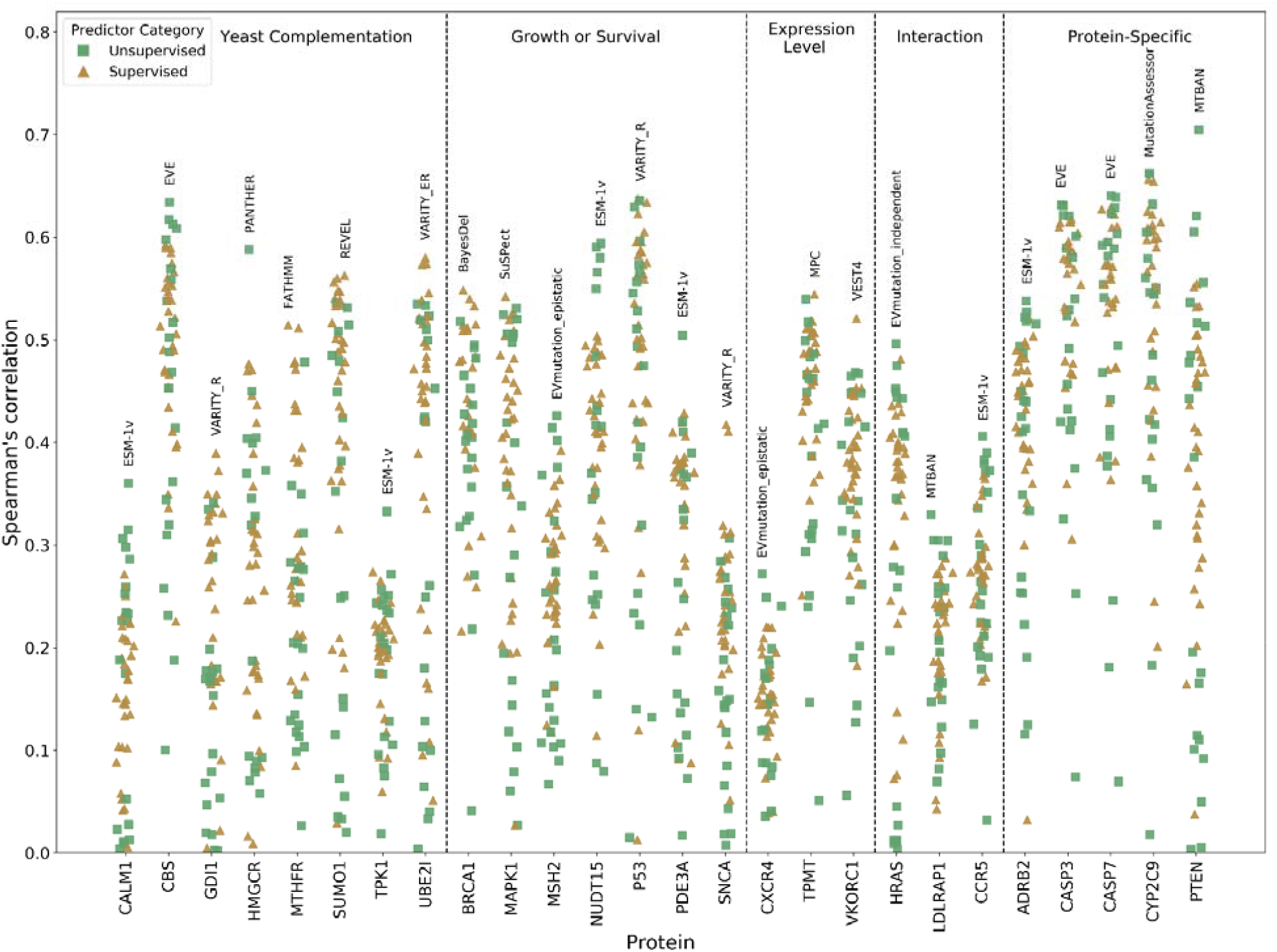
Spearman’s correlations between DMS datasets and VEPs. Spearman’s correlation between all VEPs and the selected DMS dataset for every protein. The top-performing VEP by Spearman’s correlation for each protein is labelled on the plot. DMS experiments are grouped by the type of fitness assay used where “yeast complementation” describes an assay where the human gene is used to compensate for the lack of activity in an essential yeast gene. “Growth or survival” includes any assay where growth rate is assessed apart from yeast complementation assays. “Expression level” refers to VAMP-seq or other assays that quantify the amount of protein produced. “Interaction” includes any assay involving quantifying a proteins interaction level with binding partners such as yeast two-hybrid. “Protein-specific” includes any other assay tailored to the function of a particular protein that does not easily fall into another category.

Low correlations with all VEPs were observed for several DMS datasets in our previous study, specifically for TPK1 and CALM1. The expansion of this analysis with further DMS datasets has highlighted additional cases where all VEPs fall below 0.4 Spearman’s correlation with the DMS data: CXCR4, GDI1 and LDLRAP1. Interestingly, all but one of the DMS datasets with below 0.4 Spearman’s correlation with all VEPs were carried out in yeast systems (complementation assays in CALM1, TPK1, GDI1 and a two-hybrid assay for LDLRAP1); the exception was CXCR4, which was assessed in human cells by expression level. On the other hand, some yeast assays did show high correlations, so it is likely that there are strong protein-specific factors influencing this trend. Some of the highest correlations between VEP output and DMS results observed in this study involved DMS assays that were tailored specifically to the function of the protein being assessed (‘Protein-specific assays’ in Figure 1). Other common DMS approaches such as measuring protein expression levels by VAMP-seq (Matreyek *et al*, 2018) or cell-surface expression, or measuring specific protein interaction affinities tended to be less correlated with VEP predictions or produced mixed results.

We improved upon our previous VEP rank score calculations by performing a comparison between all pairs of VEPs using the Spearman’s correlation between each VEP and DMS data across only variants for which both VEPs produced predictions. This resolves the issue of VEPs being compared across variants that are not necessarily shared between them. For example, some VEPs output predictions for every possible amino acid substitution, while others output predictions only for missense variation possible via a single-nucleotide change. Moreover, some VEPs do not output predictions across the entire lengths of all proteins. According to our methodology, VEPs receive a point for ‘winning’ each pairwise comparison, and the total score is then divided by the total number of pairwise comparisons the VEP participated in. We averaged this metric for each VEP across all DMS datasets to produce a final rank score that can be interpreted as the average proportion of VEPs that each VEP performs better than across all DMS datasets (Fig 2).

**Figure 2.**
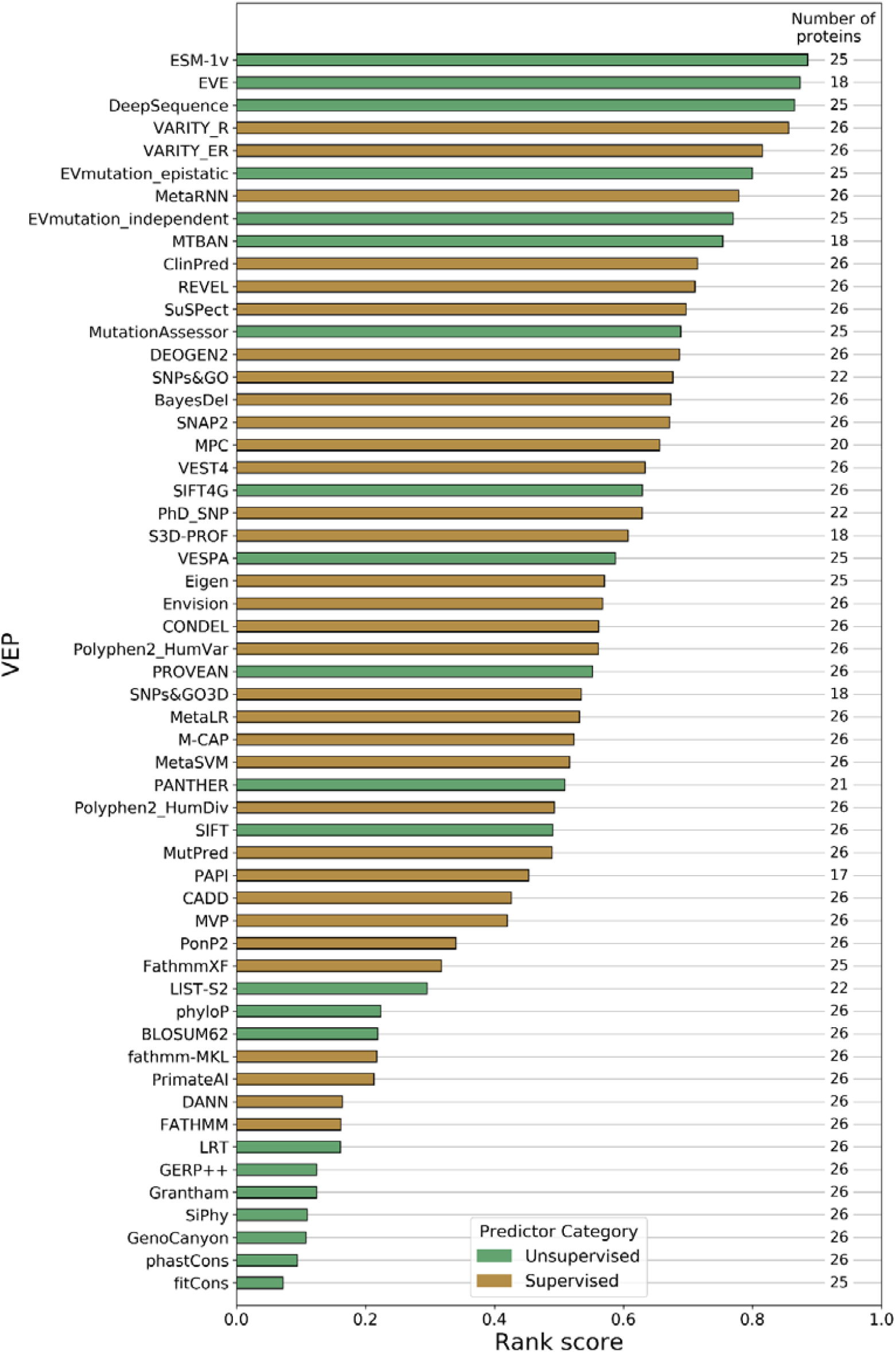
Overall ranking of VEP performance based correlation with DMS data. Rank scores for each VEP based on the Spearman’s correlation between VEP predictions and DMS data across all proteins using only shared variants by pairwise comparisons. The number of proteins for which predictions of each VEP are available are shown on the plot.

Using both the new (Fig 2, Table S5) and old (Table S6) ranking methods, the top performing VEP was ESM-1v, a new unsupervised protein language model that produces functional predictions by zero-shot inference (Meier *et al*, 2021). Like many other unsupervised predictors, ESM-1v is trained using large numbers of proteins sequences, but unlike other methods, ESM-1v is not trained using an alignment specifically related to the protein of interest. It is instead pre-trained on a large database of 98 million protein sequences. Zero-shot prediction is the application of a model to an entirely new task without any task-specific training (Lampert *et al*, 2009). Here the pre-constructed ESM-1v model is used to directly infer fitness effects for any protein with no additional training or even fine-tuning for the target proteins. In addition to performing top in our analysis, ESM-1v is considerably faster and easier to run than other top unsupervised methods (EVE, DeepSequence and EVmutation), as generating a multiple sequence alignment or training a new model for every protein is not required.

VARITY was the top-ranking supervised VEP in our analysis. VARITY is based on a gradient boosted trees model and has an innovative approach to weighting training data by predicted quality (Wu *et al*, 2021). The model gives two scores, VARITY_R which includes only rare pathogenic variants (minor allele frequency <0.5%) in the core training set and VARITY_ER which includes only very rare pathogenic variants (minor allele frequency < 1x10^−6^). It must be noted that, like Envision (Gray *et al*, 2018), VARITY uses some DMS datasets during its training, specifically ten of the same datasets we have used to assess predictors in this analysis (*UBE2I, SUMO1, CALM1, TPK1, GDI1, MTHFR, CBS, BRCA1, PTEN* and *TPMT*); therefore, data circularity may be inflating the performance estimates of VARITY. Importantly, however, after exclusion of these DMS datasets from the benchmarking analysis, VARITY_R and VARITY_ER retain 4^th^ and 5^th^ ranked places respectively, and the rank scores even improve marginally (Table S7). Thus, the strong performance of VARITY does not appear to be due to data circularity.

The other top performers, EVE (Frazer *et al*, 2021) and DeepSequence, were both developed by the same group and each makes use of an unsupervised variational autoencoder to learn the latent rules underlying a multiple sequence alignment based on the protein of interest. Performance of the two VEPs is very similar, although EVE performs slightly better. EVE scores are constrained to a range between 0 and 1 to aid with interpretability and pre-calculated results are available to download online for a subset of human proteins, while DeepSequence outputs unconstrained log likelihood ratios and does not offer any pre-calculated results.

### Performance of DMS compared to VEPs against datasets of pathogenic and benign missense variants

One of the most interesting applications of DMS data is in directly predicting the effects of clinically relevant variants. While data circularity often negatively influences our ability to determine how effective supervised VEPs are for this purpose, known clinical labels have no impact on the assessment of the predictive ability of experimentally derived, fully independent DMS data, and theoretically, a minimal impact on unsupervised VEPs. To assess the performance of DMS datasets at predicting actual clinical outcomes in comparison to unsupervised VEPs, we used known pathogenic missense variants from ClinVar (pathogenic and likely pathogenic) (Landrum *et al*, 2014) and HGMD (public) (Stenson *et al*, 2003), while putatively benign variants were obtained from gnomAD (Karczewski *et al*, 2020), excluding those that were also present in the pathogenic set. These datasets are used to calculate receiver operating characteristic area under the curve (ROC AUC) statistics for each VEP and DMS dataset. We calculated the ROC AUC for every protein with at least 10 pathogenic and 10 putatively benign missense variants. We also supplemented the *CALM1* dataset by adding variants from *CALM2* and *CALM3*, which have identical amino acid sequences, and identified additional pathogenic *SNCA* variants in the literature (Fevga *et al*, 2021; Daida *et al*, 2022; Kapasi *et al*, 2020).

Similar to our ranking analysis, we compared the ROC AUC of every pair of unsupervised predictors or DMS score sets using only variants shared between them (providing at least 10 ClinVar and gnomAD variants were shared). The method that produces the higher AUC in each pairwise comparison gains one point. Figure 3 shows the rankings of each predictor based on its mean rank across every protein. We selected the best-ranking DMS score per protein to represent the overall DMS rankings. The DMS datasets showed highly heterogeneous performance, ranking first for *TP53, SNCA*, and *BRCA1*, but performing considerably worse for *VKORC1, MAPK1, MTHFR* and *TPK1*. We note that, although DMS outperformed all VEPs for *TPK1* in our previous study, the inclusion of more pathogenic missense variants here (increasing from 8 to 14) has substantially affected its performance. Interestingly, the top-performing unsupervised VEPs from the DMS-correlation analysis (ESM-1v, EVE and DeepSequence) are also those that perform the best, on average, against the clinical data.

**Figure 3.**
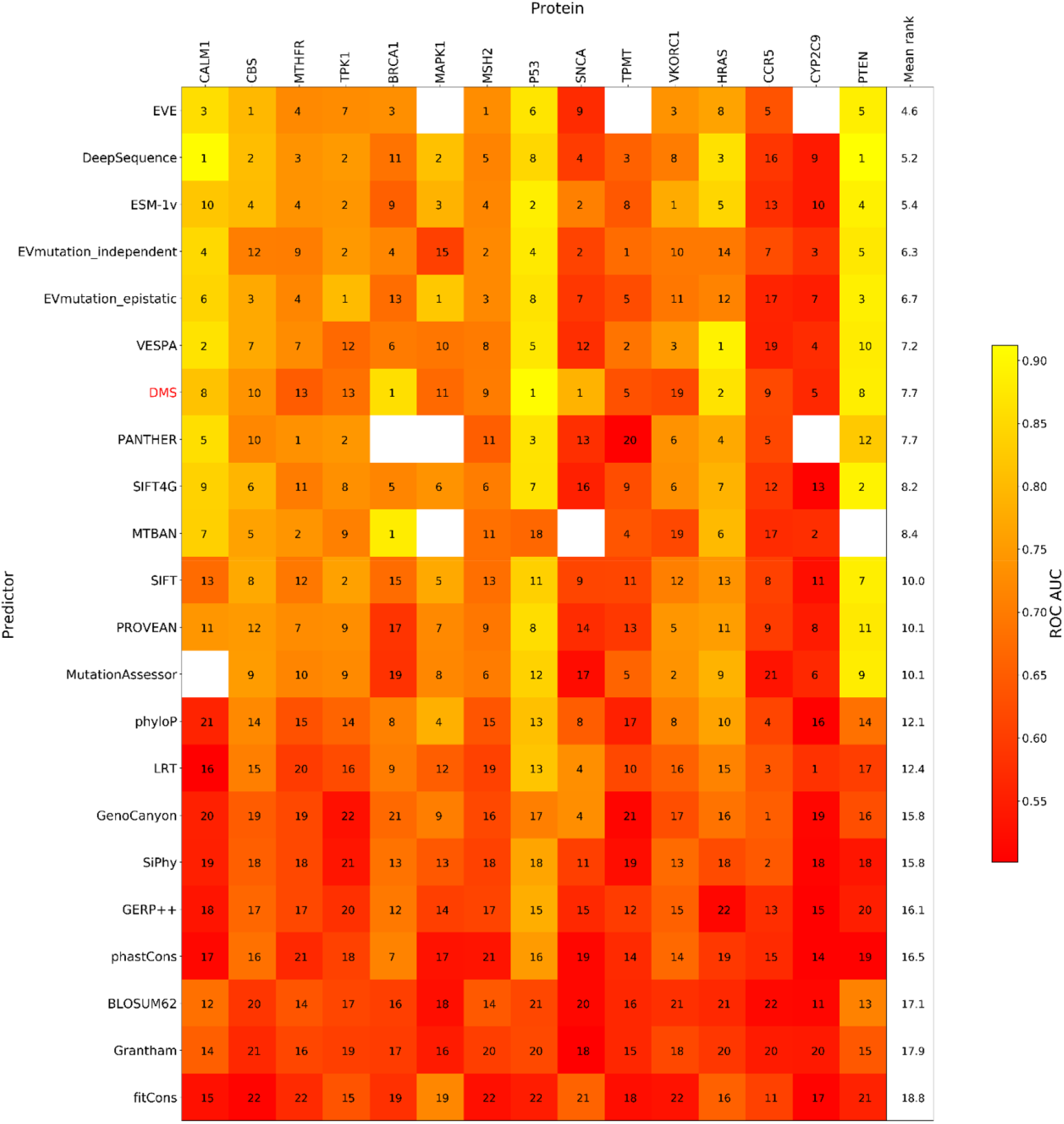
Ranking of DMS and unsupervised VEPs using clinical missense variants. The rankings of DMS and unsupervised VEPs by ROC AUC using shared variants. The colour scale of the heatmap shows the ROC AUC achieved by every predictor for classifying pathogenic and putatively benign variants, while the numbers indicate the relative ranking of all predictors for each protein. Rank ties are assigned the same rank as the top-ranking member of the group.

Consistently low ROC AUCs across all predictors, including DMS, was noted for *CYP2C9* and *CCR5*. The *CYP2C9* protein is involved in drug metabolism, and the selected DMS assay in this protein measured the variant activity on a single substrate (Amorosi *et al*, 2021). It is likely that the ClinVar and gnomAD databases are not suitable for identifying such pharmacogenetic variants, as they would appear benign until a patient was exposed to the drug. Furthermore, variant activity may alter depending on different substrates, as *CYP2C9* metabolises a large number of drug molecules. *CCR5* is a cell surface chemokine receptor and ‘pathogenic mutations’ from HGMD are typically associated with altered interaction with the human immunodeficiency virus (Howard *et al*, 1999). Similar to pharmacogenetic variants in *CYP2C9*, it is likely that a large number of these variants would be present in gnomAD, thus greatly reducing the apparent predictive performance of all methods for this protein.

For several proteins, our pathogenic and putatively benign samples are highly unbalanced (Table S8), which can potentially lead to ROC AUCs being an overly optimistic assessment of performance. The alternative is to use precision-recall curves, which focus on correct prediction of the positive (pathogenic) class but are sensitive to sample balance, making comparisons between proteins difficult. We calculated average precision scores for all predictors where the pathogenic class was less than 50% of the putatively benign class. Our results indicate that the pathogenic class is poorly predicted in *CCR5, CYP2C9, SNCA, TPK1* and *TPMT* by most predictors including DMS (Table S9).

### Benchmarking unsupervised VEPs on large clinical datasets

The issues of type 1 and 2 data circularity apply primarily to supervised VEPs; in contrast, unsupervised VEP predictions cannot be overtly influenced in the same way, as these methods are not trained using labelled data. It is still possible that some unsupervised VEPs are tweaked based on performance against clinical observations that could re-introduce type 1 circularity into performance assessments but, in general, we consider unsupervised VEPs immune to these forms of bias. As data circularity is far less likely in unsupervised VEPs, the use of traditional benchmarks with clinical data for these methods is likely to be a much better reflection of actual performance than for supervised VEPs.

To assess the performance of all unsupervised variant effect predictors against clinical data, we identified all human proteins with at least 10 pathogenic or likely pathogenic missense variants in ClinVar, and 10 other missense variants in gnomAD, leaving us with 985 proteins. Where possible, we obtained predictions from 18 unsupervised VEPs for all variants in these proteins. To compensate for the fact that some VEPs were unable to make predictions for all missense variants in a protein, we again used a pairwise ranking approach, whereby every pair of unsupervised VEPs were compared by ROC AUC. Figure 4A shows the distributions of these rank scores for each of the unsupervised predictors across all proteins.

**Figure 4.**
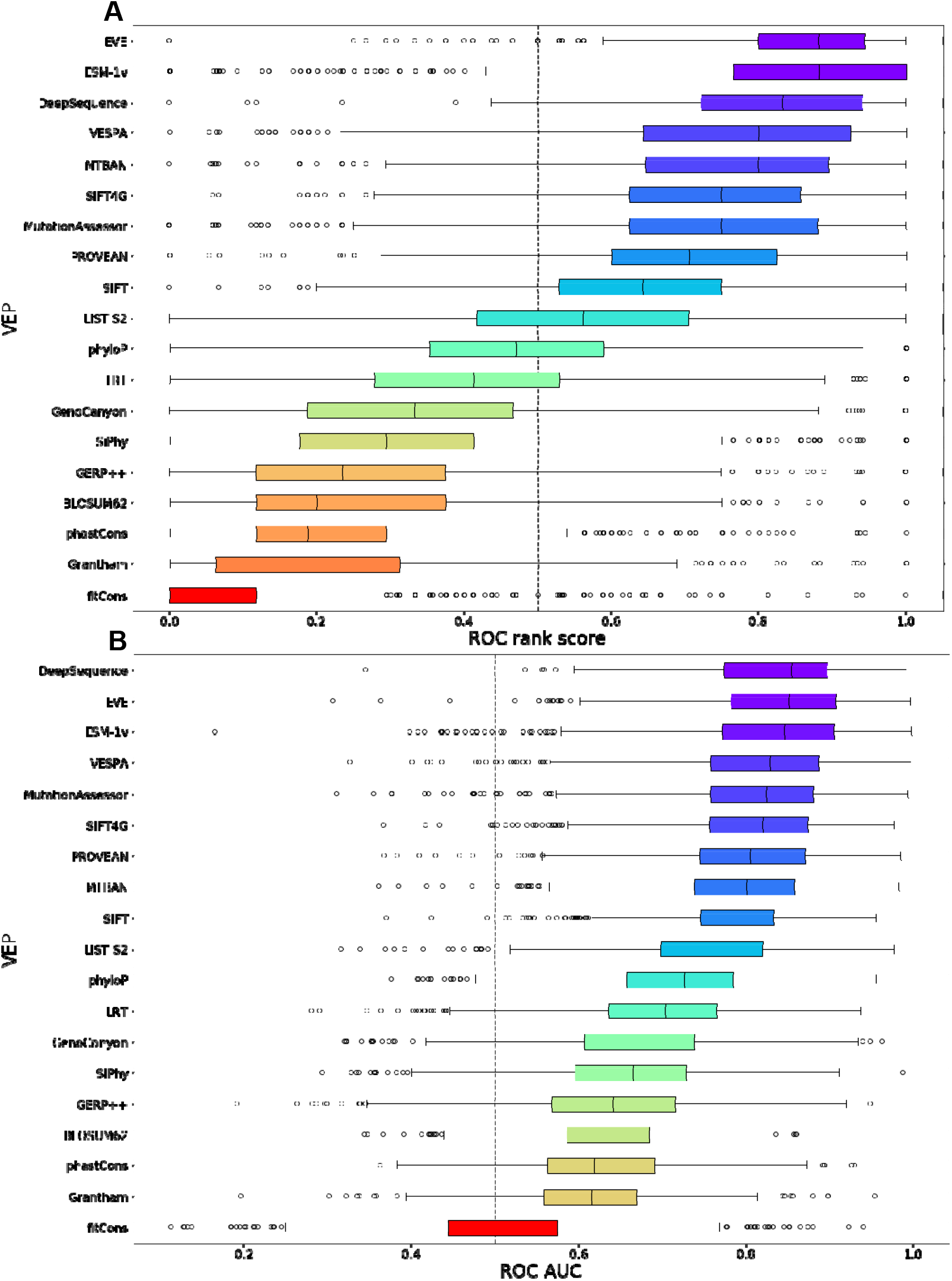
The performance of unsupervised VEPs against clinical missense variants. A) The distribution of ROC AUC-based rank scores for unsupervised VEPs on ClinVar and gnomAD variants from 985 proteins. Outliers are plotted as individual points when they occur 1.5 times the interquartile range beyond the 1^st^ or 3^rd^ quartile. An orange line indicates the median of each distribution. B) Distribution of the raw ROC AUCs for unsupervised VEPs on ClinVar and gnomAD variants from 985 proteins. Outliers are plotted as individual points when they occur 1.5 times the interquartile range beyond the 1^st^ or 3^rd^ quartile. A black line indicates the median of each distribution. EVmutation is excluded from this analysis due to predictions being available for only a very limited number of proteins.

The top-performing VEPs by median rank score were EVE and ESM-1v which, along with DeepSequence, have median ROC AUCs above 0.84 (Fig 4B). Overall, the results obtained by ranking unsupervised VEPs using clinical data were similar to their relative ranking against the DMS data. Nucleotide conservation metrics and substitution matrices are relatively poor predictors of clinical effects, while the top four VEPs are all based on advanced unsupervised machine learning methodology. It has been noted that nucleotide-based alignments (such as those that form the basis of GERP++, SiPhy and PhyloP) are considerably noisier than protein alignments (Wernersson & Pedersen, 2003) and that protein-based alignments allow for more distantly related sequences to be included in the alignment (Pearson, 2013). A combination of these factors may be contributing to the relatively poor performance of nucleotide-based conservation metrics.

Unfortunately, due to data circularity, we cannot include supervised VEPs in this analysis in a fair manner. However, given the striking correlation between our DMS rank score and the mean ROC rank score from Figure 4A (Fig 5), this strongly suggests that our findings regarding the relative performance of supervised and unsupervised VEPs against DMS data are a valid representation of likely predictor performance against clinically relevant human missense variants.

**Figure 5.**
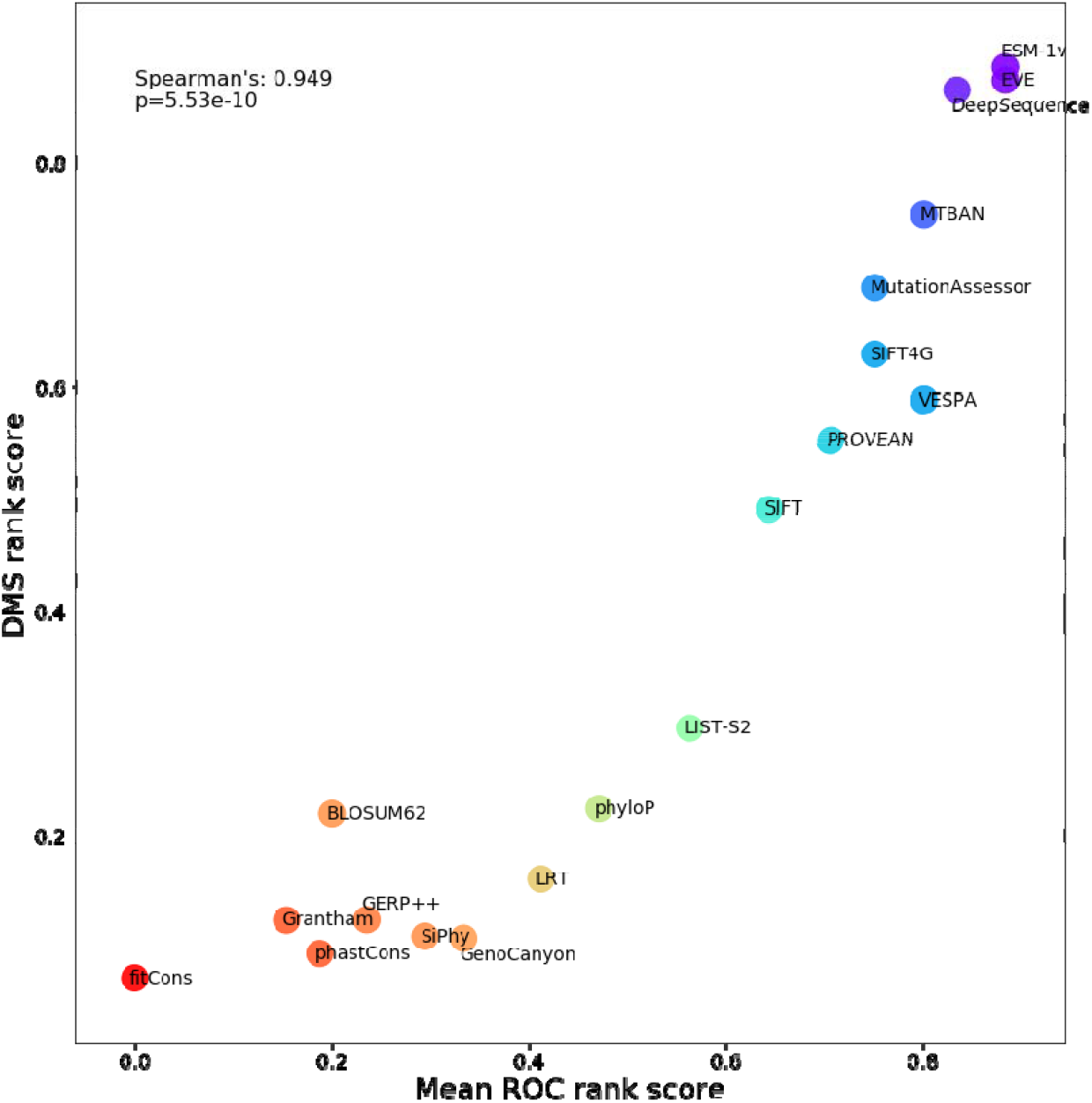
The relationship between VEP rank score and ROC rank score for unsupervised VEPs. The rank score of unsupervised VEPs from Figure 2 plotted against the ROC rank score from Figure 4A. The identity of each unsupervised VEP is indicated on the chart.

## Discussion

Our updated analysis produced some interesting results in terms of predictor ranking; DeepSequence remained a highly ranked method, but was joined by ESM-1v, EVE and VARITY. The presence of many new predictors among the most highly ranked indicates that VEP methodology is continuing to improve. With the exception of DeepSequence, supervised VEPs were previously superior to most unsupervised methods. Our present results indicate that most of the top-10 VEPs are now unsupervised, demonstrating that multiple unsupervised methodologies are viable for VEP development, and that researchers are taking the potential for bias seriously and making efforts to avoid introducing it into new VEPs. No particular machine learning technique is dominant among top-ranking VEPs, indicating that multiple approaches to variant effect prediction remain powerful with their unique advantages and disadvantages.

The excellent performance of ESM-1v in the correlation analysis is particularly interesting, not due to its nature as an unsupervised VEP, but because it had no access to a protein-specific multiple sequence alignment like EVE, DeepSequence and EVmutation. While type 1 and 2 data circularity poses no issue for these predictors, sampling bias from the database used to construct MSAs still has the potential to influence predictions in some proteins. ESM-1v has demonstrated that even this source of bias can be mitigated, although not entirely eliminated, as language models are still trained using a sequence database, albeit a very large and varied one. We used ESM-1v in a zero-shot setting, where no MSA generation took place, but it is also possible to use the model in a “few-shot” setting, where a protein-specific MSA is provided to assist with protein-specific predictions. The authors of the method found that using the model in a few-shot context improved predictions slightly (Meier *et al*, 2021), but we were unable to successfully run this model on our system.

For the supervised methods Envision and VARITY, this analysis does not constitute a truly independent benchmark, as some of the DMS datasets from our benchmark were also a part of their training data. VARITY in particular may be somewhat optimised for predicting the results of DMS experiments in general, but its strong performance on datasets that were not used in its training suggests that this is not a major issue. It seems likely that more newly developed VEPs will incorporate DMS data in the future. While it makes little sense to exclude DMS datasets as a potential source of training data, it does mean that future benchmarking using this data may carry the same caveats as benchmarking traditional supervised predictors using variant databases. Similar scenarios will likely arise for any new source of benchmarking data, as it will eventually be used as a training dataset for new VEPs. We must continue to be vigilant regarding the data used to train VEPs, and where possible ensure that fully independently derived data is used for benchmarking.

Our ROC analysis included nine further proteins over our previous study and made use of numerous additional variants deposited in ClinVar since 2018, and variants not previously included from HGMD. While DMS data did not perform as the top predictor for the majority of proteins, it was still often among the top methods. Notably, DMS ranked first for three proteins, which was more than any individual VEP. However, DMS also performed quite poorly more often than the top VEPs, demonstrating that DMS datasets are highly heterogeneous in their performance in disease variant classification. We previously claimed that DMS experiments based on growth rate tended to be more representative of human disease mutations compared to those based on protein expression levels or other assays. With an expanded set of DMS data and additional variants, this conclusion no longer seems valid, as some DMS assays based on expression levels and quantifying protein-protein interactions predicted disease as well as those based on yeast complementation or general growth rate. It is crucial that we learn what factors make a DMS dataset reliable for this purpose, whether they be related to the target protein specifically, the choice of experimental assays, or other technical issues with the experiments. Is there some way we can predict *a priori* whether a DMS dataset will be predictive of variant pathogenicity? Interestingly, there is little correspondence between the median VEP correlation with DMS datasets, and the performance of DMS datasets for variant classification (Fig S1A).

However, it is notable that the three most clinically predictive datasets were all for tumour suppressors (*P53, BRCA1* and *PTEN*), which also have fairly high correlations with VEPs. It may be that the observed growth rate changes in these DMS studies are more reflective of the actual functional changes seen in human disease than for other classes of genes. In addition, there was a significant correlation between the relative ranking of DMS among predictors for classification of clinical variants and correlation with DMS predictions (Fig S1B). Those DMS studies that do correlate well with VEPs tend to perform more highly when compared against them on clinical data.

The *CALM1* dataset was an outlier with high ROC AUC (and good prediction of the positive class) but low correlation with all VEPs. Despite the low correlation, numerous VEPs are still more predictive of clinical outcome than the DMS data (Fig 3). It may be that the VEPs and DMS are picking up different disease-related features in the *CALM* genes that are uncorrelated but still lead to decent performance against clinical data.

The biggest outlier in the opposite direction with good DMS correlation and poor clinical performance was *CYP2C9* (Fig S1A). Both *NUDT15* and *CYP2C9* are proteins involved in drug metabolism that contain pharmacogenetic variants influencing the activity and specificity of those proteins. Pharmacogenetic variants may have no impact on phenotype until an individual is treated with a particular drug; thus, databases of pathogenic mutations are greatly depleted in such variants. In addition, the lack of notable phenotype also means that many pharmacogenetic variants may be included in databases of benign variants, and in gnomAD. While VEPs achieved moderately strong correlations with DMS for *CYP2C9*, indicating that the measured fitness values were being reasonably well predicted by the VEPs, predictions of clinical data in *CYP2C9* was poor across all methods. This is likely due to the influence of pharmacogenetic variants being present in gnomAD, thus reducing the apparent predictive ability of all VEPs and the DMS data.

Our results continue to indicate that benchmarking using independent variant effect datasets is a powerful strategy for reducing data circularity when assessing VEP performance. The potential of DMS for direct variant effect prediction remains exciting, although care should be taken to ensure that the assay used is indicative of phenotypic outcome. With less than three years of additional data, we more than doubled the number of human DMS datasets in this analysis, and it is likely that with projects like the Atlas of Variant Effects (www.varianteffect.org), the availability of such datasets, and their utility for protein variant interpretation, will explode.

## Methods

### DMS identification and criteria

We retained 13 DMS datasets in human proteins from our previous analysis and identified a further 19 studies with publically available datasets from MAVEDB (Esposito *et al*, 2019) (https://www.mavedb.org/) and literature searches. We applied a threshold of 5% minimum coverage of all amino acid variants within the target protein to prevent any particularly low-coverage studies from skewing our results. This prevented a *SCN5A* dataset being included (Glazer *et al*, 2020). We also excluded datasets for *NCS1* and *TECR* obtained from MAVEDB as no methodology was published with them.

### VEP score retrieval

Most VEP predictions were retrieved from the dbNSFP database version 4.2 (academic) (Liu *et al*, 2020). Scores were retrieved for the transcript that matches the canonical Uniprot sequence for each protein. As dbNSFP is a nucleotide-resolution database, there are instances where multiple nucleotide variants map to the same amino acid substitution. In these cases the mean of the VEP scores mapping to the same substitution were taken.

SIFT was run locally using the UniRef90 database (Suzek *et al*, 2015) to generate multiple sequence alignments.

EVmutation scores were obtained from the EVcouplings pipeline (mutation stage). We used the Uniref100 database to generate alignments and default settings as found in: https://github.com/debbiemarkslab/EVcouplings/blob/develop/config/sample_config_monomer.txt except changing the minimum_column_coverage setting to 20 to reduce large alignment gaps.

DeepSequence was run locally using alignments generated by the EVcouplings pipeline with default settings. EVE results were partially retrieved online from: https://evemodel.org/ and others were run locally using default settings on a GPU.

ESM-1v results were obtained by adapting the example at: https://github.com/facebookresearch/esm/blob/main/examples/variant-prediction/predict.py and running locally on a GPU. The final score is the mean of esm1v_t33_560_UR90S_1, esm1v_t33_560_UR90S_2, esm1v_t33_560_UR90S_3, esm1v_t33_560_UR90S_4 and esm1v_t33_560_UR90S_5 outputs.

Sources for all VEPs can be found in Table S3.

### Correlation analysis

For each protein we had DMS data for, we selected a single DMS dataset to be representative of it in our analysis. We selected the dataset with the highest median Spearman’s correlation to all VEP predictions for that protein to help prevent outliers from influencing the choice of set.

Spearman’s correlation was calculated using the scipy.stats.spearmanr() function of the python scipy package.

### Rank score calculation

For each selected DMS dataset, every pair of VEPs was compared to each other using only variants shared between both of them and the DMS data. The Spearman’s correlation of both VEPs to the DMS data was calculated and the ‘winning’ predictor was awarded 1 point. The ‘losing’ predictor was awarded 0 points while a tie resulted in 0.5 points each. Once every pair of VEPs was compared, the final score was divided by the number of times each VEP was tested to determine the proportion of all other VEPs that it ‘beat’. The mean of this value was taken across all proteins which had predictions for that particular VEP to generate the final rank score.

### Variant identification

For calculation of ROC and PR AUC values, we used the gnomAD database version 2.1.1 (Karczewski *et al*, 2020) (https://gnomad.broadinstitute.org/) as a source of putatively benign variants. While these variants certainly contain some recessive and low-penetrance pathogenic variants, gnomAD filters out individuals with severe paediatric disease and their first-degree relatives. This means that gnomAD should be depleted in pathogenic variants relative to the population and serves as a useful estimate of benign variation.

Pathogenic variants were primarily obtained from the ClinVar database of clinically relevant variants (August 2022 update) (Landrum *et al*, 2014) (https://www.ncbi.nlm.nih.gov/clinvar/) and also from HGMD (free version) (Stenson *et al*, 2003) (https://www.hgmd.cf.ac.uk/ac/index.php). Additional pathogenic variants from *SNCA* were found though a literature search (Fevga *et al*, 2021; Daida *et al*, 2022; Kapasi *et al*, 2020). Any pathogenic variants present in the gnomAD dataset were removed from the gnomAD set.

Variants in *CALM2* and *CALM3* were used to supplement *CALM1* variants as all three proteins share the same primary structure, although they differ at the genomic level.

### ROC calculation

ROC AUC values were calculated using the sklearn.metrics.roc_auc_score() function of the sklearn python package. Pathogenic variants were labelled as true positives and gnomAD variants were labelled as true negatives. In the case of VEPs with inverted score scales (such as SIFT) where lower values indicate increased likelihood of pathogenicity, the positive and negative classes were switched before the ROC AUC was calculated.

ROC scores were calculated in a series of pairwise comparisons between predictors using only shared variants. The predictor with the highest AUC received 1 point and the ‘loser’ received 0 points. When all predictors for a protein had been compared, they were ranked based on the total number of points received.

### PR calculation

A summary of the precision-recall cuve was calculated using the sklearn.metrics.average_precision_score() function of the sklearn python package. Average precision score is a statistic that summarises PR curves as a weighted mean of precisions over each threshold (Turpin and Scholer, 2006).

## Data Availability

A compiled dataset of all DMS scores and VEP predictions used to perform this analysis is available at: https://doi.org/10.6084/m9.figshare.21581823.v1.

## Acknowledgements

This project was supported by the European Research Council (ERC) under the European Union’s Horizon 2020 research and innovation programme (grant agreement No. 101001169). JM is a Lister Institute Research Fellow. BL was supported by the MRC Precision Medicine Doctoral Training Programme.

## Supplemental Material

**Figure S1.**
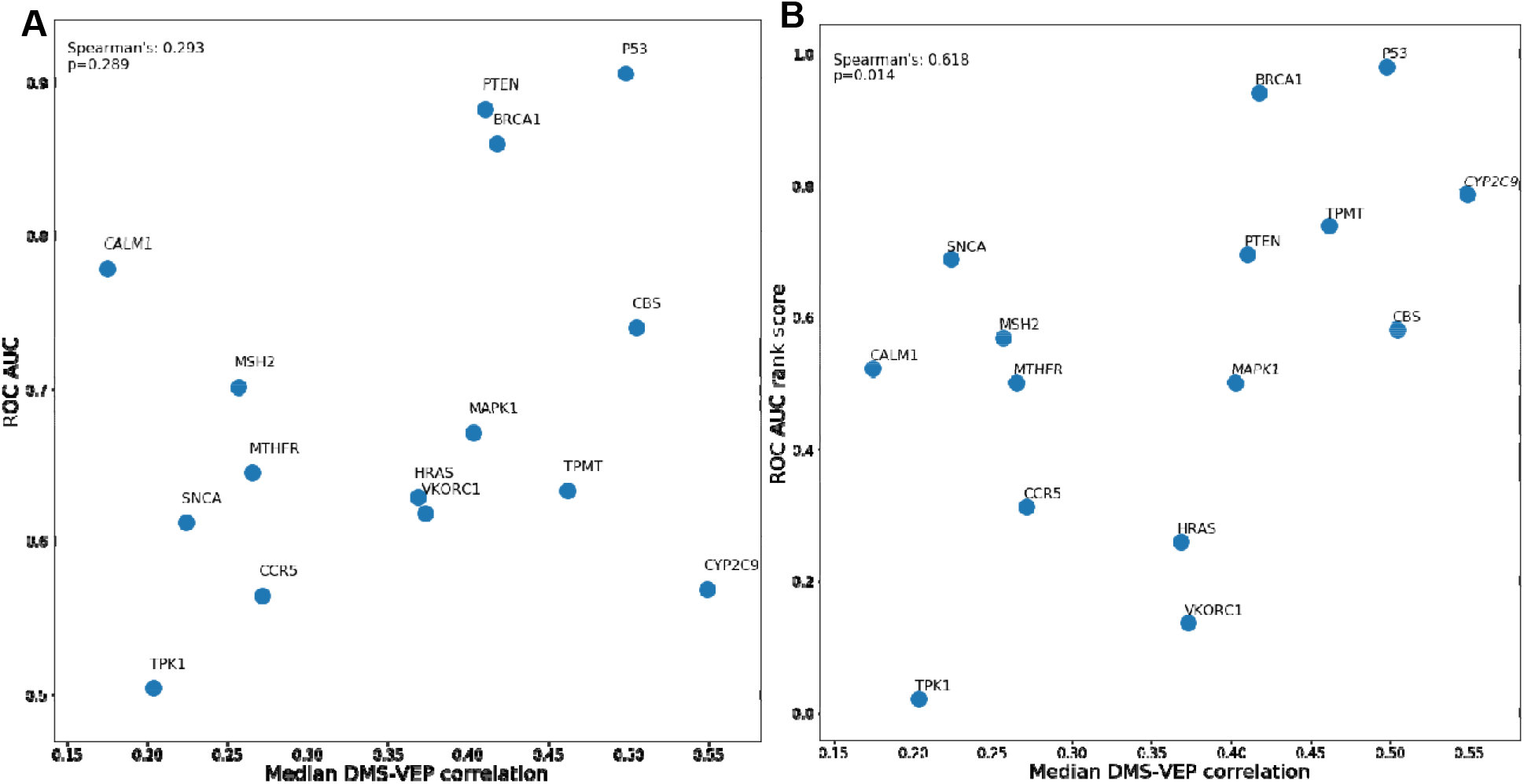
The relationship between median VEP-DMS correlation and ROC AUC. **A)** The median correlation between each DMS dataset and VEP (from Figure 1) plotted against the ROC AUC of each DMS study, separating ClinVar and HGMD pathogenic variants from putatively benign gnomAD variants. **B)** The median correlation between each DMS dataset and VEP (from Figure 1) plotted against the ROC AUC-based rank score of each DMS study, separating ClinVar and HGMD pathogenic variants from putatively benign gnomAD variants.

**Table S1 - Summary of all DMS assays**. Details of all DMS assays used in this analysis including the source and selected score set for the correlation-based analyses.

**Table S2 -** Mapping between assay names assigned by the authors and internal identifiers used in figures and tables in this analysis

**Table S3 - Correlation between DMS assays**. The Spearman’s correlations and p-values between all DMS replicates and alternative assays in the dataset.

**Table S4 - Details of all VEPs assessed**. Source, references and classifications of all VEPs analysed in this study.

**Table S5** - **Rank score for VEPs across all DMS datasets**. Individual correlation-based rank scores calculated for each VEP across all proteins and the final rank score averaged across all proteins.

**Table S6 - Old rank score calculations**. Rank scores as defined in (Livesey & Marsh, 2020). The rank score represents the mean, normalised Spearman’s correlations across all proteins.

**Table S7 - Rank scores without the VARITY training data**. Correlation-based rank scores calculated using a subset of proteins that excludes the training data of VARITY.

**Table S8 - The numbers of pathogenic and putatively benign variants used to calculate ROC AUC**. The number of pathogenic variants includes data from both ClinVar and HGMD. A minimum of 10 variants in both classes was required for inclusion in Figure 3.

**Table S9 - Average precision scores of VEPs for proteins with imbalanced data**. Average precision scores for all VEPs and DMS datasets across nine proteins where the pathogenic dataset was less than half the size of the putatively benign set.

