## Supplementary material for "Updated benchmarking of variant effect predictors using deep mutational scanning": Table S1

**Table S1. Summary of all DMS assays**

| **Protein (Uniprot)** | **Functional assay** | **Selected score (internal name)** | **Percentage coverage of all amino acid variants (%)** | **Reference** |
| --- | --- | --- | --- | --- |
| *CALM1* (P0DP23) | Competitive growth assay in yeast | DMS_flipped | 64.04 | (Weile *et al*, 2017) |
| *TPK1* (Q9H3S4) |  | DMS_flipped | 68.90 |  |
| *SUMO1* (P63165) |  | DMS_flipped | 88.59 |  |
| *UBE2I* (P63279) |  | DMS_flipped | 85.38 |  |
| *CBS* (P35520) | Competitive growth assay in yeast | DMS_refined_lowB6 | 64.41 | (Sun *et al*, 2020) |
| *GDI1* (P31150) | Competitive growth assay in yeast | DMS | 51.40 | (Silverstein *et al*, 2021) |
| *HMGCR* (P04035) | Competitive growth assay in yeast | DMS_no_statin | 99.89 | (Jiang, 2019) |
| *LDLRAP1* (Q5SW96) | Yeast two-hybrid | DMS_OBFC1 | 99.03 |  |
| *MTHFR* (P42898) | Competitive growth assay in yeast | DMS_25_A222V | 99.85 | (Weile *et al*, 2021) |
| *BRCA1*(a) (P38398) | Yeast two-hybrid and phage display. | N/A | N/A | (Starita *et al*, 2015) |
| *BRCA1*(b) (P38398) | Growth rate of HAP1 cells | DMS_b | 5.19 | (Findlay *et al*, 2018) |
| *MAPK1* (P28482) | Growth rate of A375 cells | DMS_DOX | 99.56 | (Brenan *et al*, 2016) |
| *MSH2* (P43246) | Rescue of MMR-deficient HAP1 cells | DMS | 94.38 | (Jia *et al*, 2021) |
| *NUDT15* (Q9NV35) | Drug resistance assay. | DMS_sensitivity | 94.16 | (Suiter *et al*, 2020) |
| *P53*(a) (P04637) | Growth assay in the presence of P53 agonists | N/A | N/A | (Giacomelli *et al*, 2018) |
| *P53*(b) (P04637) | Growth rate assay in human cells | DMS_b | 39.37 | (Kotler *et al*, 2018) |
| *PDE3A* (Q14436) | DNMDP sensitivity in a glioblastoma cell line | DMS_DNMDP_100 | 36.41 | (Garvie *et al*, 2021) |
| *SNCA* (P37840) | Yeast growth rate hindered by toxic aggregates (reverse survival) | DMS_Miconazole | 97.26 | (Newberry *et al*, 2020) |
| *CCR5* (P51681) | Antibody binding | DMS_Ab2D7 | 97.97 | (Heredia *et al*, 2018) |
| *CXCR4* (P61073) | Surface expression levels in human cells | DMS_expression | 99.36 |  |
| *TPMT* (P51580) | Protein stability assessed by FACs (VAMP-seq) | DMS | 79.25 | (Matreyek *et al*, 2018) |
| *PTEN*(a) (P60484) |  | N/A | N/A |  |
| *PTEN*(b) (P60484) | Disruption of an artificial gene circuit in yeast | DMS_highqual_b | 85.73 | (Mighell *et al*, 2018) |
| *VKORC1* (Q9BQB6) | Protein stability assessed by FACs (VAMP-seq) | DMS_VAMP | 87.02 | (Chiasson *et al*, 2020) |
| *HRAS* (P01112) | Yeast two-hybrid | DMS_g12v | 87.30 | (Bandaru *et al*, 2017) |
| *ADRB2* (P07550) | Reporter expression | DMS_0.65 | 99.40 | (Jones *et al*, 2020) |
| *CASP3* (P42574) | Apoptotic activity assessed by a microfluidic system. | DMS | 28.63 | (Roychowdhury & Romero, 2022) |
| *CASP7* (P55210) |  | DMS | 29.17 |  |
| *CYP2C9* (P11712) | Activity profiling by click-seq | DMS_activity | 65.97 | (Amorosi *et al*, 2021) |
