## Supplementary material for "Updated benchmarking of variant effect predictors using deep mutational scanning": Table S4

**Table S3. Details of all VEPs assessed**

| **VEP or score** | **Classification** | **Data source** | **Reference** |
| --- | --- | --- | --- |
| SIFT | Unsupervised | Run locally, scripts available from: <https://sift.bii.a-star.edu.sg/www/code.html> | (Sim *et al*, 2012) |
| SIFT4G | Unsupervised | dbNSFP 4.2 | (Vaser *et al*, 2016) |
| phyloP | Unsupervised | dbNSFP 4.2 | (Pollard *et al*, 2010) |
| BLOSUM62 | Unsupervised | <https://www.ncbi.nlm.nih.gov/Class/FieldGuide/BLOSUM62.txt> | (Henikoff & Henikoff, 1992) |
| LRT | Unsupervised | dbNSFP 4.2 | (Chun & Fay, 2009) |
| SiPHy | Unsupervised | dbNSFP 4.2 | (Garber *et al*, 2009) |
| GERP++ | Unsupervised | dbNSFP 4.2 | (Davydov *et al*, 2010) |
| Grantham | Unsupervised | Available from referenced paper | (Grantham, 1974) |
| EVmutation (independent and epistatic) | Unsupervised | Run locally, available from: <https://github.com/debbiemarkslab/EVcouplings> | (Hopf *et al*, 2017) |
| LIST_S2 | Unsupervised | dbNSFP 4.2 | (Malhis *et al*, 2020) |
| PROVEAN | Unsupervised | dbNSFP 4.2 | (Choi *et al*, 2012) |
| PANTHER | Unsupervised | <https://snps.biofold.org/snps-and-go/snps-and-go.html> | (Thomas & Kejariwal) |
| MutationAssessor | Unsupervised | dbNSFP 4.2 | (Reva *et al*, 2011) |
| DeepSequence | Unsupervised | Run locally, scripts available from: | (Riesselman *et al*, 2018) |
| GenoCanyon | Unsupervised | dbNSFP 4.2 | (Lu *et al*, 2015) |
| EVE | Unsupervised | Downloaded from <https://evemodel.org/> or run locally using scripts at: <https://github.com/OATML/EVE> | (Frazer *et al*, 2021) |
| MTBAN | Unsupervised | <http://mtban.kaist.ac.kr/> | (Kim *et al*, 2021) |
| VESPA | Unsupervised | <https://zenodo.org/record/5905863> | (Marquet *et al*, 2021) |
| ESM-1v | Unsupervised | Run locally, scripts available from: <https://github.com/facebookresearch/esm> | (Meier *et al*, 2021) |
| phastCons | Unsupervised | dbNSFP 4.2 | (Siepel & Haussler, 2005) |
| fitCons | Unsupervised | dbNSFP 4.2 | (Gulko *et al*, 2015) |
| Polyphen-2 (HumVar and HumDiv models) | Supervised | dbNSFP 4.2 | (Adzhubei *et al*, 2010) |
| Eigen | Supervised^1^ | dbNSFP 4.2 | (Ionita-Laza *et al*, 2016) |
| Envision | Supervised | <https://envision.gs.washington.edu/shiny/envision_new/> | (Gray *et al*, 2018) |
| SuSPect | Supervised | <http://www.sbg.bio.ic.ac.uk/suspect/index.html> | (Yates *et al*, 2014) |
| DEOGEN2 | Supervised | dbNSFP 4.2 | (Raimondi *et al*, 2017) |
| SNPs&GO | Supervised | <https://snps.biofold.org/snps-and-go/snps-and-go.html> | (Capriotti *et al*, 2013) |
| MPC | Supervised | dbNSFP 4.2 | (Samocha *et al*, 2017) |
| VEST4 | Supervised | dbNSFP 4.2 | (Carter *et al*, 2013) |
| SNAP2 | Supervised | <https://rostlab.org/services/snap2web/> | (Hecht *et al*, 2015) |
| PhD_SNP | Supervised | <https://snps.biofold.org/snps-and-go/snps-and-go.html> | (Capriotti *et al*, 2006) |
| S3D-PROF | Supervised | <https://snps.biofold.org/snps-and-go/snps-and-go-3d.html> | (Capriotti & Altman, 2011) |
| SNPs&GO3D | Supervised | <https://snps.biofold.org/snps-and-go/snps-and-go-3d.html> | (Capriotti & Altman, 2011) |
| MutPred | Supervised | dbNSFP 4.2 | (Pejaver *et al*, 2020) |
| PonP2 | Supervised | <http://structure.bmc.lu.se/PON-P2/> | (Niroula *et al*, 2015) |
| PonPS | Supervised | <http://structure.bmc.lu.se/PON-PS/> | (Niroula & Vihinen, 2017) |
| Fathmm-XF | Supervised | dbNSFP 4.2 | (Rogers *et al*, 2018) |
| Fathmm | Supervised | dbNSFP 4.2 | (Shihab *et al*, 2013) |
| Fathmm-MKL | Supervised | dbNSFP 4.2 | (Shihab *et al*, 2015) |
| PrimateAI | Supervised | dbNSFP 4.2 | (Sundaram *et al*, 2018) |
| VARITY (R and ER models) | Supervised | <http://varity.varianteffect.org/> | (Wu *et al*, 2021) |
| REVEL | Supervised | dbNSFP 4.2 | (Ioannidis *et al*, 2016) |
| CONDEL | Supervised | <http://bbglab.irbbarcelona.org/fannsdb/> | (González-Pérez & López-Bigas, 2011) |
| MetaLR | Supervised | dbNSFP 4.2 | (Dong *et al*, 2015) |
| MetaSVM | Supervised | dbNSFP 4.2 | (Dong *et al*, 2015) |
| M-CAP | Supervised | dbNSFP 4.2 | (Jagadeesh *et al*, 2016) |
| PAPI | Supervised | <http://papi.unipv.it/> | (Limongelli *et al*, 2015) |
| MVP | Supervised | dbNSFP 4.2 | (Qi *et al*, 2021) |
| CADD | Supervised | dbNSFP 4.2 | (Kircher *et al*, 2014) |
| DANN | Supervised | dbNSFP 4.2 | (Quang *et al*, 2015) |
| MetaRNN | Supervised | dbNSFP 4.2 | (Li *et al*, 2021) |
| ClinPred | Supervised | dbNSFP 4.2 | (Alirezaie *et al*, 2018) |
| BayesDel | Supervised | dbNSFP 4.2 | (Feng, 2017) |

^1^ Eigen is trained using an unsupervised spectral method, but includes PolyPhen-2 as a feature, which may introduce data circularity into its assessment. It has thus been labelled as supervised.
